# TransMetaSegmentation (TMS): a transcriptome-based segmentation method for spatial metabolomic data

**DOI:** 10.1101/2024.06.12.598521

**Authors:** Yongheng Wang, Kayle J. Bender, Weidi Zhang, Siyu Lin, Elizabeth K. Neumann, Aijun Wang

## Abstract

Matrix-assisted laser desorption/ionization (MALDI) mass spectrometry imaging (MSI) is a powerful analytical tool that enables the visualization and comparison of relative abundances of metabolites across samples, shedding light on biological processes and disease mechanisms. Techniques such as scSpatMet enable the determination of cell boundaries and cell types through staining with 35 cell marker antibodies. Yet, distinguishing subpopulations of cells, such as astrocytes, oligodendrocytes, and neuronal clusters in the brain, remains challenging using antibodies. In this context, we introduce TransMetaSegmentation (TMS), an alternative segmentation and cell typing method that integrates MALDI MSI imagery with single-cell spatial transcriptomic analysis. This approach not only delineates cell boundaries and defines cell types based on a number of marker genes but also effectively allocates metabolites to specific cell types in a high-throughput manner. We anticipate that TMS will improve the granularity of MALDI MSI analyses, advance our understanding of metabolic alterations in diseases, and have an impact on various fields within biomedical sciences.

## Introduction

Mass spectrometry imaging (MSI) has been employed to uncover the intricate distribution patterns of a vast array of molecular species, including metabolites, lipids, peptides, proteins, and glycans, with high spatial precision^1^. It offers invaluable insights into the complex interplay of biomolecules and their roles in physiological processes, disease mechanisms, and therapeutic interventions. Principal ionization methods employed in MSI include Secondary Ion Mass Spectrometry (SIMS), Desorption Electrospray Ionization (DESI), and Matrix-Assisted Laser Desorption/Ionization (MALDI)^2^. SIMS can characterize hydrophobic molecules from 0 to 1000 Daltons, offering spatial resolutions ranging from 100 nm to 1 μm^3^. DESI facilitates the identification of metabolites with molecular masses ranging from 0 to 2000 Daltons, at spatial resolutions of 40 to 200 μm^4^. MALDI is capable of detecting metabolites across a broad molecular weight spectrum, from 0 to 100,000 Daltons, with spatial resolutions approximately from 10 to 100 μm^5, 6^. Due to its broad applications, MALDI MSI has been utilized for extensive metabolic profiling in complex tissues. However, a limitation of MALDI MSI analysis is the difficulty in defining the cell boundaries, which hinders its ability to determine cell-specific metabolic profiles and unveil cellular functions and interactions within complex tissues.

To address this challenge, some antibody-based methods have been developed. For example, MALDI-immunohistochemistry (MALDI-IHC) uses photocleavable mass tags for antibody labeling and demonstrates that it can detect up to 12 different biomarkers simultaneously^7^. These cell marker antibodies can divide the tissue into smaller areas based on the presence or absence of specific antibody staining, although it does not provide single-cell resolution^7^. Another technology, scSpatMet^8^, offers a finer resolution by integrating two advanced methods: multiplex imaging mass cytometry (IMC)^9^ which labels 35 antibodies with different metal tags, and 3D-SMF^10^, a metabolic imaging technique. Consequently, scSpatMet allows for the delineation of cell boundaries and cell types and facilitates the assignment of more than 200 metabolites to various cell types. However, antibody-based methods have their drawbacks. They sometimes fail to segment every cell, leaving gaps in cell type information for unstained cells. Moreover, these methods can be restrictive when defining cell types based only on a limited set of proteins, lacking the precision provided by transcriptomic profiling.

In addition to IMC or proteomic analysis, single-cell spatial transcriptomic analysis represents an alternative way to define thousands of cell types and subclusters in a high-throughput manner^11, 12^. Single-cell spatial transcriptomics technologies can quantify the transcripts of thousands of genes with a resolution close to 200 nm. One such technology is Xenium^13^, which determines cell boundaries and cell types through different steps. It utilizes DAPI staining along with CellPose^14, 15^ and other methods to accurately define cell borders. Additionally, Xenium characterizes cell types based on the expression of specific marker genes within each cell. This is achieved through a library of nucleic acid probes, each tailored with a sequence that is complementary to the RNA transcript of a targeted gene. The probes hybridize with their corresponding transcripts, allowing for the characterization of gene expression across different cells^13^.

## Result

We developed a transcriptomic-based segmentation technique for metabolomic analysis. Specifically, we prepared two serial sections of the brain: one for MALDI MSI and the other for Xenium analysis. By integrating the information from both MALDI MSI and Xenium, we can assign metabolites to different cell types in a high throughput manner (Fig. 1). Using MALDI MSI, we putatively identified more than 200 lipids in mouse brains, with mass ranging from 747.7491 to 985.5282 m/z, at a high spatial resolution (10 µm x 10 µm)^16^. These lipids span several categories, including glycerolipids (triacylglycerols and trialkylglycerols), glycerophospholipids (glycerophosphocholines [PC], glycerophosphates [PA], glycerophosphoserines [PS], glycerophosphoethanolamines [PE], glycerophosphoglycerols [PG]), and sphingolipids (notably hexosylceramides [HexCer] and sulfatides [SHexCer]) (Fig. 2). MALDI MSI data unveils elevated levels of certain lipids in a Canavan mouse brain such as PC(44:10) (Fig. 2d), HexCer(38:0;O4) (Fig. 2m), and HexCer(40:0;O2) (Fig. 2n), alongside reduced levels of others like PC(32:4) (Fig. 2b), PC(35:4) (Fig. 2e), and HexCer(39:1;O3) (Fig. 2l). While MALDI MSI provides insights into the upregulation and downregulation of lipids, it cannot identify the cell types involved in these metabolic changes. To delineate the cell boundaries and determine the cell types, we employed a spatial single-cell technology, Xenium (Fig. 3). Xenium enabled the identification of various cell types including astrocytes, endothelial cells, perivascular macrophages, and vascular leptomeningeal cells within the hippocampus region (Fig. 3a-3e). In addition, Xenium reveals the different patterns of gene expression in Canavan and the control brains (Fig. 3e-3m).

**Fig. 1.**
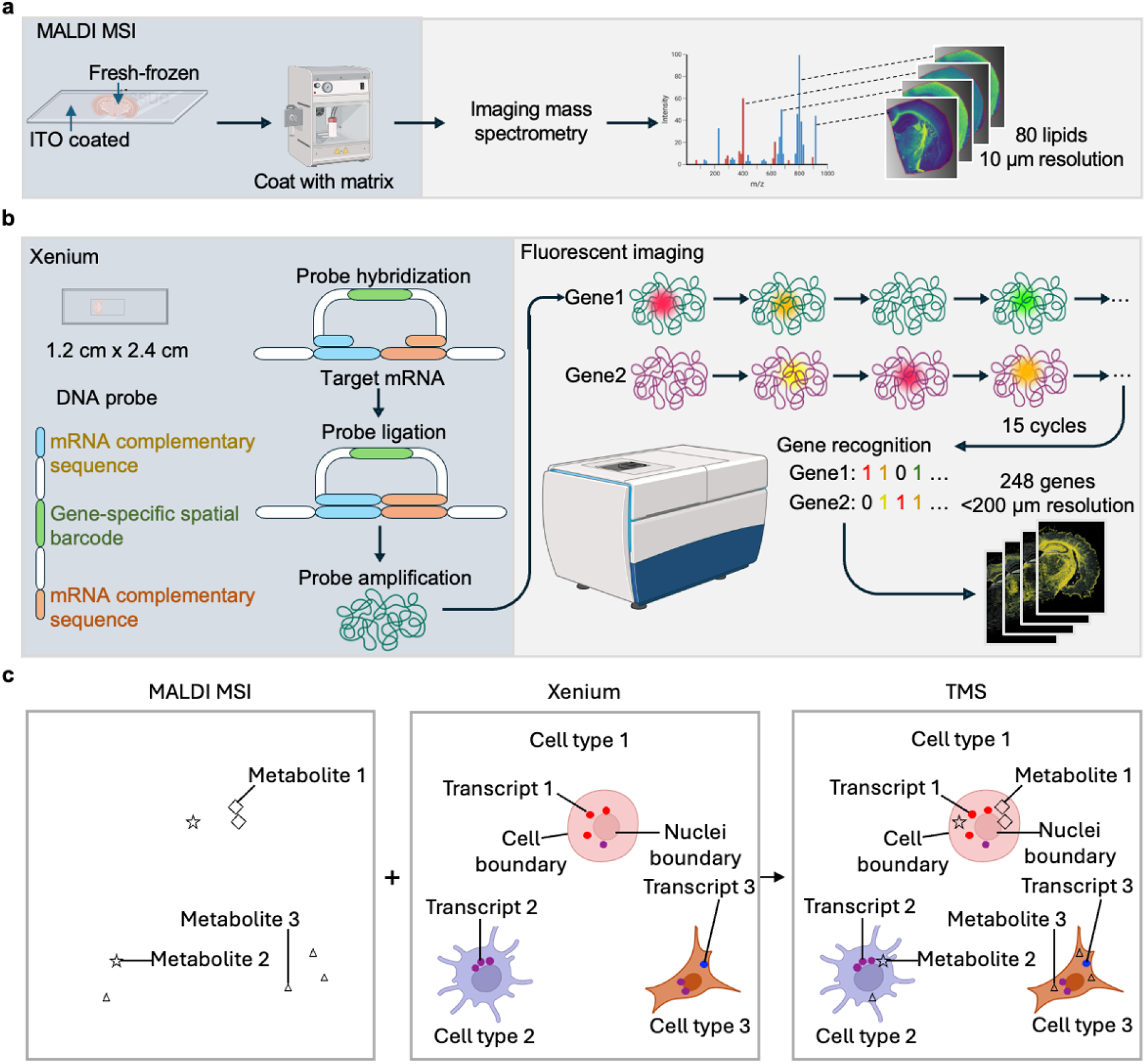
Schematic of TMS. **a**, Procedure for MALDI MSI. **b**, Xenium workflow. **c**, TMS: integration of the information from MALDI MSI and Xenium.

**Fig. 2.**
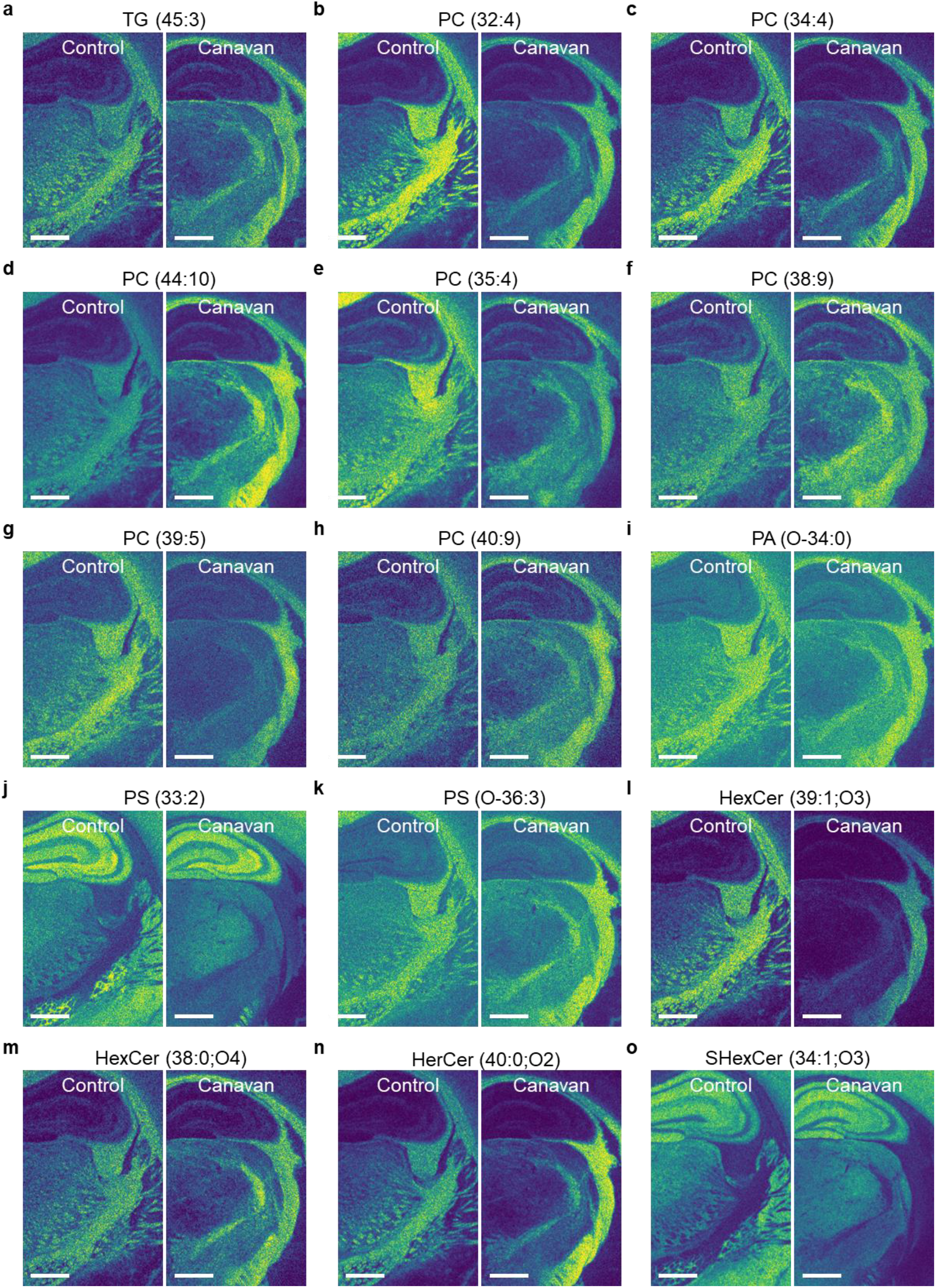
Spatial distribution of glycerolipids, glycerophospholipids, and sphingolipids. **a**, Spatial distribution of triacylglycerols TG(45:3) (m/z=797.6205). **b-d**, Spatial distribution of glycerophospholipids or glycerophosphocholines PC(32:4) (m/z= 726.5141, **b**), PC(34:4) (m/z=754.5435, **c**), and PC(44:10) (m/z=904.5819, **d**). **e-h**, Spatial distribution of diacylglycerophosphocholines PC(35:4) (m/z=806.5113, **e**), PC(38:9) (m/z=822.5077, F), PC(39:5) (m/z=860.5596, G), and PC(40:9) (m/z=850.5369, **h**). (**i**) Spatial distribution of glycerophosphate PA(O-34:0) (m/z=701.4839). **j**,**k**, Spatial distribution of glycerophosphoserines PS (33:2) (m/z=746.4831, **j**) and PS(O-36:3) (m/z=772.5548, **k**). **l-n**, Spatial distribution of sphingolipids hexosyl ceramides HexCer(39:1;O3) (m/z=808.6352, **l**), HexCer(38:0;O4) (m/z=812.6275, **m**), and HerCer(40:0;O2) (m/z=824.6284, **n**), **o**, Spatial distribution of sphingolipid sulfatides Sulfoglycosphingolipids SHexCer(34:1;O3) (m/z=834.4918). Scale bars are 800 μm.

**Fig. 3.**
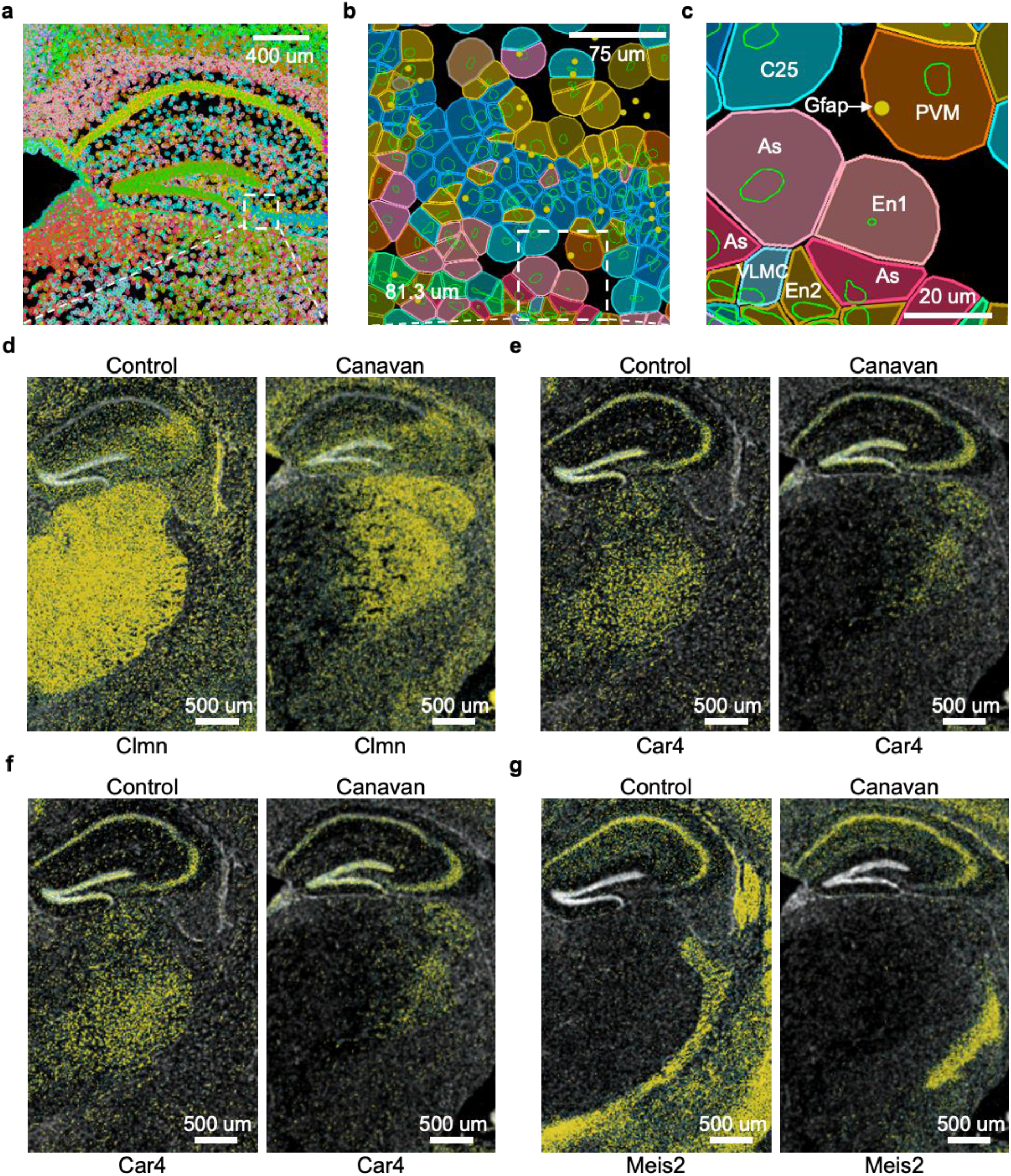
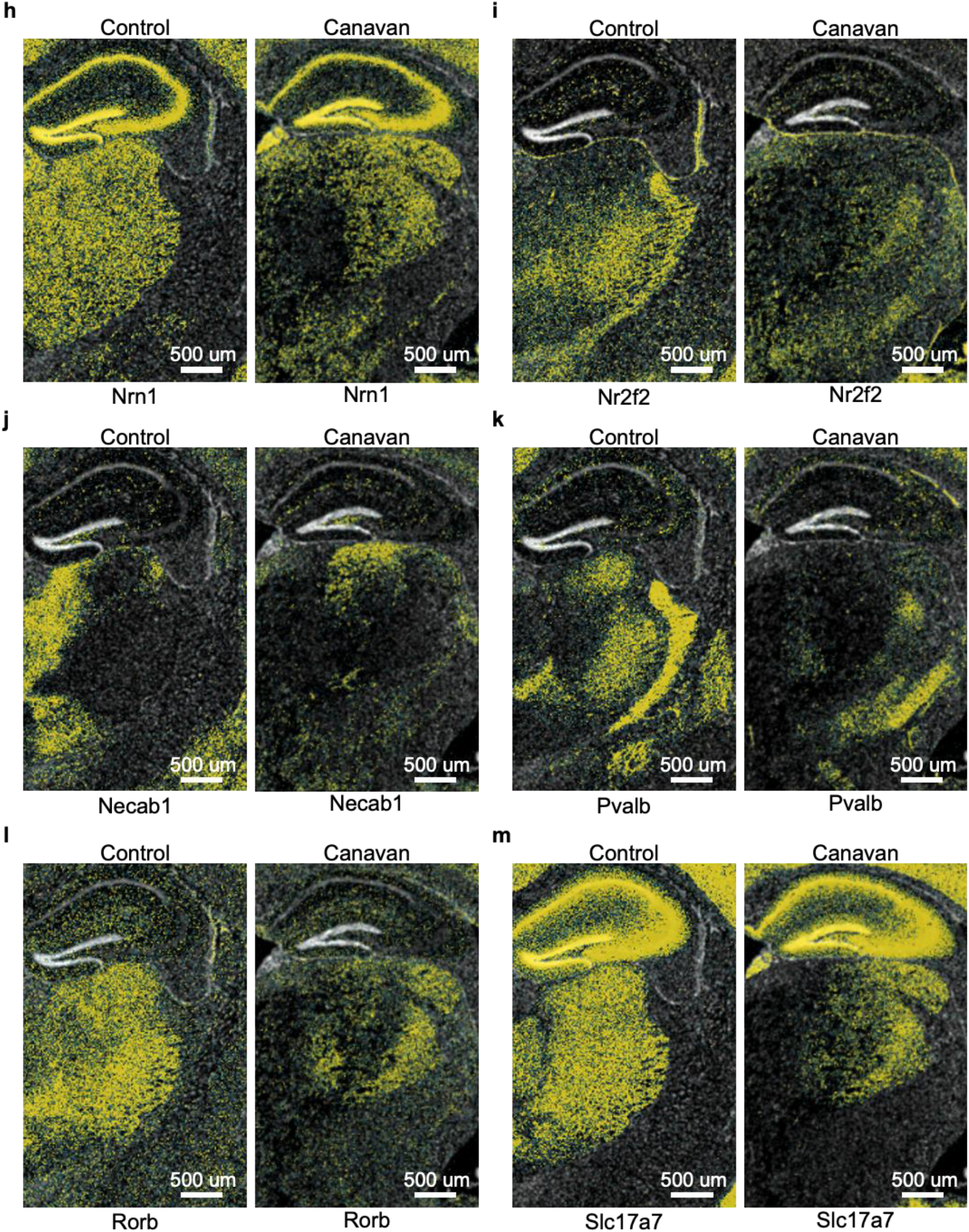
Xenium profiles the cell types and boundaries. **a**-**c**, Cell boundaries are defined by nucleus-staining and CellPose. Cell types are determined by gene expression patterns. Xenium data visualization within Xenium Explorer presents a hippocampus with cells displayed as colored dots (**a**), sorted by cell type (**b**). A detailed regional view shows nucleus areas bounded by green lines (**c**). **d-m**, Differential expression of genes Clmn (**d**), Car4 (**e**), Car4 (**f**), Meis2 (**g**), Nrn1 (**h**), Nr2f2 (**i**), Necab1 (**j**), Pvalb (**k**), Rorb (**l**), and Slc17a7 (**m**). As, astrocyte; PVM, perivascular macrophages; VLMS, vascular leptomeningeal cells; En, endothelial cell.

Using TMS, we attribute the metabolites to various cell types. This enabled us to compare the lipid levels between the same cell types in control and Canavan brains (Fig. 4-5). Notably, in Canavan disease, both oligodendrocytes and astrocytes exhibited elevated levels of SHexCer(42:2;O3) (Fig. 4a-4f) and PG(44:10) (Fig. 5g-5l), along with reduced levels of PC(41:5) (Fig. 4g-4l). Additionally, astrocytes exhibit elevated PS(O-34:4) (Fig. 5e).

**Fig. 4.**
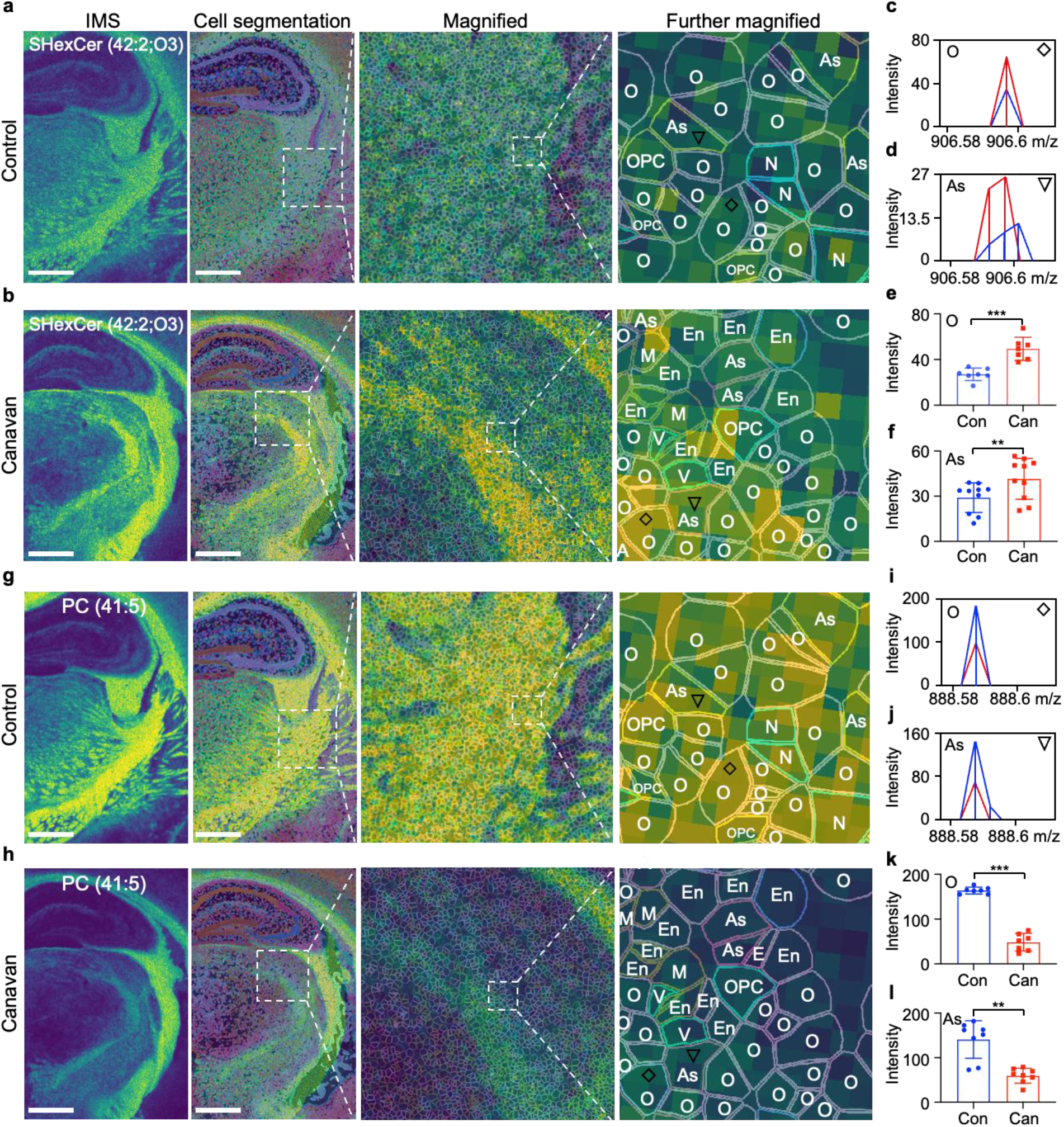
TMS enables the evaluation of the lipids in oligodendrocytes and astrocytes. **a**-**f**, Overlay an image from MALDI MSI with a cell segmentation image from Xenium yields a single-cell spatial lipidomic image. Spatial distribution (**a, b**) and quantification (**c-f**) of SHexCer(42:2;O3) (m/z=906.5968) in oligodendrocytes (rhombus) and astrocytes (triangle). **g-l**, Spatial distribution (**g, h**) and quantification (**i-l**) of PC(41:5) (m/z=888.5872) in oligodendrocytes (star) and astrocytes (triangle). As, astrocyte; O, oligodendrocyte; En, endothelial cell; M, microglia; N, neuron; OPC, oligodendrocyte progenitor cells; P, pericyte; V, vascular leptomeningeal cells. Scale bars are 800 µm. Values are presented as mean ± SD. p values determined by Mann-Whitney U tests. ***p < 0.001. *p-value < 0.01. *p < 0.05.

**Fig. 5.**
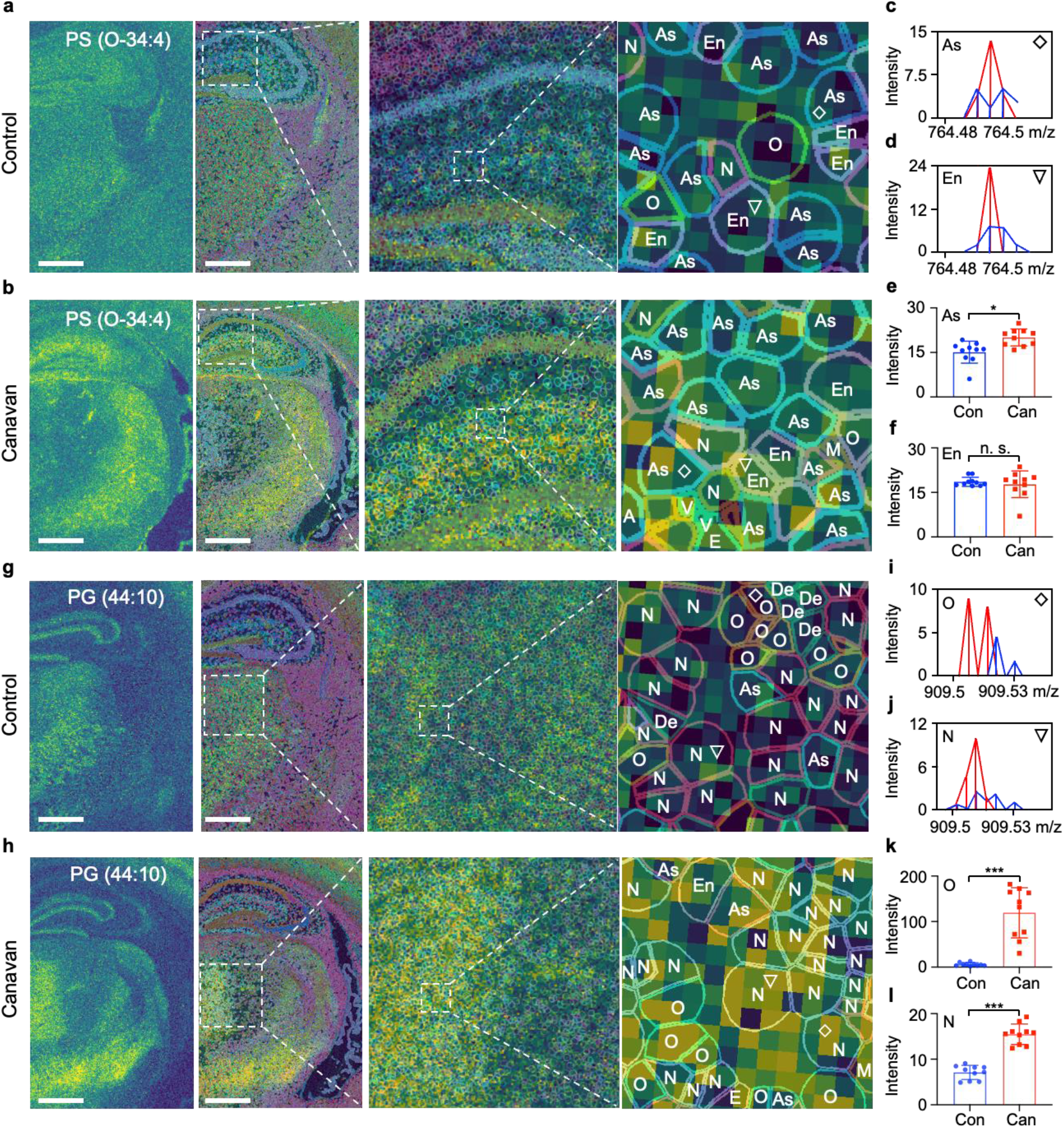
TMS allows quantification of lipids in different cell types. **a**-**f**, Spatial distribution (**a, b**) and quantification (**c-f**) of PS(O-34:4) (m/z=764.4892) in astrocytes (rhombus) and endothelial cells (triangle). **g-l**, Spatial distribution (**g, h**) and quantification (**i-l**) of PG(44:10) (m/z=909.5117) in oligodendrocytes (star) and neurons (triangle). As, astrocyte; De, dendritic cell; O, oligodendrocyte; En, endothelial cell; N, neuron. Scale bars are 800 µm. Values are presented as mean ± SD. p values determined by Mann-Whitney U tests. ***p-value < 0.001. **p-value < 0.01. *p-value < 0.05. n.s., no significant difference.

## Discussion

Using MALDI MSI alone, we revealed prominent dysregulation in lipid composition, including elevated levels of certain SHexCer, PG, and PS species, and reduced levels of various PC species in Canavan brains. These findings underscore the profound metabolic disruptions associated with Canavan disease. Unfortunately, MALDI MSI requires further experimentation to specify the exact cell types responsible for these alterations. The integration of Xenium with MALDI MSI via our TMS method addresses this limitation by providing precise cell boundary information and cell type identification. This approach allows us to map the distribution of specific lipids within individual cells, revealing elevated levels of some SHexCer species in oligodendrocytes and astrocytes, and increased abundance of some PG lipid species in oligodendrocytes and neurons in Canavan brains. While this study focused on using Canavan disease and Xenium as illustrative examples, other single-cell spatial technologies such as CosMx, MERSCOPE, and Stereo-seq can also be combined with MALDI MSI data to investigate the metabolic changes in various pathological conditions.

There are notable limitations when using TMS for defining cell boundaries and determining cell types. The segmentation methods employed in spatial transcriptomic technologies typically combine DAPI staining and CellPose. While useful, this does not achieve the accuracy of antibody staining for delineating cell boundaries, particularly in non-spherical cells like star-shaped astrocytes and bipolar neurons. Additionally, relying solely on mRNA levels to identify cell types may not always correlate accurately with protein levels. Despite these challenges, TMS may present advantages in certain scenarios. TMS excels in analyzing spherical cells, as commonly found in the spleen, and can effectively differentiate among thousands of neuronal sub-clusters in the brain. In conclusion, our approach, while acknowledging the inherent limitations, leverages the strengths of TMS for detailed metabolomic analysis in complex biological systems. TMS stands out as a powerful tool for advancing our understanding of complex diseases and is poised to make a significant impact in the field of biomedical sciences.

## Materials and Method

### Animal experiments

All animal procedures were approved by The University of California, Davis, Institutional Animal Care and Use Committee (IACUC). Canavan Mice (Strain #:008607) were purchased from Jackson laboratory.

### MALDI MSI

Mouse brains were dissected, then placed on a foil paper boat, which was then filled with pulverized dry ice. Isopentane was poured over the dry ice, ensuring rapid temperature reduction and efficient freezing of the tissue to preserve its molecular and structural integrity. Once frozen, the brain was embedded in 2.6% carboxymethyl cellulose (CMC; EMD Millipore Corp., Burlington, MA), sectioned into 10 µmthick sections using a cryomicrotome (Leica Biosystems, Wetzlar, Germany), and placed on an indium tin oxide (ITO) coated slide. Subsequently, a 1,5-diaminonaphthalene (DAN, Tokyo Chemical Industry Company Ltd., Tokyo, Japan) matrix (20 mg/mL in tetrahydrofuran) was applied to the tissue using an HTX M3+ sprayer (HTX Technologies, LLC, Chapel Hill, NC) to enhance laser energy absorption and ionization of tissue molecules for mass spectrometry. The spraying conditions were set to a nozzle temperature of 40 °C, nitrogen gas pressure of 15 psi, a solvent flow rate of 50 μL/min, and a track spacing of 2 mm, with a total of five passes over the tissue. Analysis of the tissue was conducted with a timsTOF fleX dual source mass spectrometer (Bruker Scientific, Billerica, MA), operated in positive ion mode. The analysis used a raster width of 10 μm × 10 μm. To ionize the tissue molecules, 150 laser shots were fired in a single burst. The instrument settings included a global attenuator value of 0%, a local laser power of 88%, and a mass-to-charge (m/z) ratio range of 50-2000. Data analysis was carried out using SCiLS Lab version 2023 software (Bruker Scientific), and lipid identification was achieved by comparing the mass spectrometry data with known lipid profiles in the LIPID MAPS database^17^.

### Xenium

Fresh-frozen samples were sliced into 10-micrometer-thick sections and placed onto the Xenium slides. Sections were digested to make the mRNA accessible. The following reagents were added: control probes for non-specific binding assessment, genomic DNA controls for signal source assessment, and 313 probes with two target RNA complementary sequences and one gene-specific barcode. Probes with a concentration of 10 nM were hybridized to the RNA at 50°C overnight. The tissue was washed to remove the unhybridized probes. Hybridized probes were ligated at 37°C for two hours. The primers for Rolling Circle Amplification (RCA) were added to prepare for probe amplification. The probes were amplified in a two-step enzymatic process—first for one hour at 4°C and then for two hours at 37°C. Multiple copies of the gene-specific barcodes were created, increasing the signal-to-noise ratio. A second wash was performed to chemically quench the background fluorescence. The tissue sections were placed into an imaging cassette and loaded into the Xenium Analyzer for image acquisition. Afterwards, fluorescent oligonucleotides were added to the sample. During the 15 cycles of probe hybridization and imaging, Z-stacks images were captured at 0.75 μm intervals for each cycle. A spatial map of the RNA transcripts was created by stitching the Z-stacks images together, using the DAPI image as a reference. Cell segmentation was performed by using DAPI images to identify cell nuclei and CellPose.

## Acknowledgements

This work was funded by a grant from the National Institute of Health (1R21NS133881-01 to A. W.). Fig.1 was created with BioRender.com. We thank 10X Genomics for their support with the Xenium experiment.

## Author contributions

Y.W. conceived the TMS idea with guidance from A.W. and E.K.N. K.J.B. conducted the MALDI MSI experiment and provided support for MSI analysis. W.Z. assisted with TMS data analysis. S.L. helped create Fig. 3. Y.W., S.L., and W.Z. wrote the first draft. All authors reviewed the manuscript.

